# Dynamic configurations of meiotic hotspot determinants

**DOI:** 10.1101/2020.06.26.167775

**Authors:** Yu-Chien Chuang, Gerald R. Smith

## Abstract

During meiosis, appropriate DNA double-strand break (DSB) and crossover distributions are required for proper homologous chromosome segregation in most species. Linear element proteins (LinEs) of *Schizosaccharomyces pombe* are DSB hotspot determinants. Clusters of LinE-bound hotspots form within ∼200 kb chromosomal regions independent of DSB formation. Previous reports showed that LinEs form chromatin-bound, dot-like nuclear foci in nuclear spreads and in fixed cells. Here, we investigated the regulation of LinE configuration and distribution in live cells using super-resolution fluorescence microscopy. In live cells at optimal meiotic temperature (∼25°C), LinEs made long linear forms, not previously reported, in both zygotic and azygotic meiosis and shared other characteristics with the synaptonemal complex in other species. LinE structures appeared around the time of replication, underwent a dotty-to-linear-to-dotty configurational transition, and disassembled before the first meiotic division. DSB formation and repair did not detectably influence LinE structure formation, but failure of DSB formation delayed LinE structure disassembly. Several LinE missense mutations formed dotty but not linear LinE configurations. Our study reveals a second, important configuration of LinEs, which suggests that LinE complexes are involved in regulating meiotic events, such as DSB repair, in addition to their established role in DSB formation.

## Introduction

Meiosis is a special type of cell division that allows diploid cells to reduce the chromosome number by half to generate haploid gametes. In meiotic prophase, homologous chromosomes pair and undergo meiotic recombination to exchange genetic material. The physical exchange of portions of homologous chromosomes – formation of a crossover (CO) – generates recombinant chromosomes that increase the genetic diversity of the gametes. COs also form the physical connections (chiasmata) between homologous chromosomes that provide tension to ensure that homologous chromosomes segregate properly from each other in the first meiotic nuclear division (MI) (Zickler and Kleckner, 1999; Petronczki et al., 2003; Marston and Amon, 2004). Meiotic recombination is initiated by programmed DNA double-strand breaks (DSBs). The topoisomerase-like protein Spo11, first identified in budding yeast, contains the active site for meiotic DSB formation in multiple experimental organisms (Keeney et al., 1997; McKim and Hayashi-Hagihara, 1998; Dernburg et al., 1998; Baudat et al., 2000; Cervantes et al., 2000; Bowring et al., 2006; Hartung et al., 2007). In the few species examined, meiotic DSBs are non-randomly distributed and concentrated in short DNA intervals called DSB hotspots (Pan et al., 2011;Fowler et al., 2014; Lange et al., 2016).

DSB hotspot determination is well-documented in the fission yeast *Schizosaccharomyces pombe*. Linear element proteins (LinEs; Rec10, Rec25, Rec27 and Mug20) are DSB hotspot determinants (Fowler et al., 2013). Rec10 is distributed nearly uniformly along the chromosomes, while the other three LinEs co-localize at most if not all DSB hotspots with high specificity. Rec10 is necessary for all DSB formation, and the other three LinEs are required for DSB formation at nearly all DSB hotspots by stabilizing or activating the DSB-forming complex containing Rec12 (Spo11 homolog in *S. pombe*) (Martín-Castellanos et al., 2005; Davis et al., 2008; Fowler et al., 2013). The strong correlation between the distribution of LinE binding and DSB hotspots, and the requirement for LinEs for DSB formation at most hotspots, makes *S. pombe* one of the best understood models in DSB hotspot-designation studies.

The distance between two crossovers is highly regulated. To ensure proper chromosome segregation in MI, both COs and sister chromatid cohesion are needed to generate tension between homologous chromosomes. However, too-close COs may result in no cohesion between homologs, which can lead to mis-segregation (Zickler and Kleckner, 1999). Crossover interference, the occurrence of one crossover reducing the frequency of a second nearby, was found over a hundred years ago and avoids too-close COs (Sturtevant, 1915; Muller, 1916). Several models of CO interference have been proposed, but the molecular basis of most of these models is not clear (King and Mortimer, 1990; Foss et al., 1993; Foss and Stahl, 1995; Fujitani et al., 2002; Kleckner et al., 2004; Hultén, 2011). Based on both genetic and molecular evidence, Fowler *et al.* (2018) proposed a “clustering model” in which CO interference is the outcome of DSB interference, which has an identified molecular basis.

The clustering model proposes that DSB hotspots bound by LinEs along a limited chromosomal region form a compact, three-dimensional cluster before DSB formation, and a limited number of DSBs (perhaps only one) is formed per cluster (Fowler et al., 2018; Nambiar et al., 2019). With a modification of the genome-wide chromatin conformation capture (3C) technique (Dekker et al., 2002; Fowler et al., 2018), Fowler et al. (2018) showed that the DSB-determining LinEs within a roughly 200 kb region form hotspot clusters. They showed that both DSB competition and DSB interference are also limited to ∼200 kb regions. Both DSB interference and crossover interference depend on the Tel1 DNA damage-response protein kinase, consistent with crossover interference resulting from DSB interference. The physical distance of 200 kb is about 35 cM in *S. pombe* (Fowler et al., 2018), which is similar to the extent, in genetic distance, of crossover interference in *Drosophila melanogaster* and *Neurospora crassa* (Foss et al., 1993). The clustering model predicts that LinE clusters regulate crossover interference within ∼200 kb regions in *S. pombe*, as observed. However, the mechanism of formation of LinE complexes and DSB hotspot clusters is not yet known.

LinEs are distantly related to the synaptonemal complex (SC) proteins of other species (Colaiácovo et al., 2003; Lorenz et al., 2004; Loidl, 2006; Fowler et al., 2013), but *S. pombe* is considered not to form SC in meiosis. Electron microscopic studies of nuclear spreads show that LinEs form much shorter structures than the *S. cerevisiae* SC, which form more nearly continuous, end-to-end chromosomal structures (Dresser and Giroux, 1988; Bähler et al., 1993). Similarly, fluorescence microscopy of LinEs in nuclear spreads shows that LinEs form dots or short linear structures (Lorenz et al., 2004; Davis et al., 2008), and LinEs in fixed cells show dot-like foci (Davis et al., 2008; Estreicher et al., 2012; Fowler et al., 2013). However, most of these studies were done with fixed cells, which were cultured at high temperature (34°C). By contrast the cohesin subunit Rec8 shows dot-like structures in spread nuclei and fixed cells at 34°C (Davis et al., 2008; Fowler et al., 2013) but intact, end-to-end chromosomal structures in live cells at 26°C using super-resolution microscopy (Ding et al., 2016). This result indicates, first, that Rec8 forms different configurations under different culture conditions and, second, that *S. pombe* chromosomes may form large but more fragile SC-like structures that partially disassemble upon nuclear spreading or fixation. Based on these findings, it is important to investigate whether LinEs show end-to-end, SC-like chromosomal structures in live cells at 25°C.

We studied LinE structures in live cells using super-resolution fluorescence microscopy. LinEs formed linear structures at 25°C in both zygotic and azygotic meiosis (described below), and the linear structure showed liquid-crystalline properties like the SC in other species (Rog et al., 2017). The LinE linear structures were also found in synchronized azygotic meiosis at 25°C but not at 34°C, and they underwent a dot-to-line-to-dot transition, showing that there are at least two different LinE configurations. Moreover, we investigated the genetic requirements for intact LinE focus formation and factors that trigger its disassembly. Our results reveal a new form of LinE-chromatin structure and suggest that LinEs have a function in addition to DSB hotspot-determination and CO interference regulation.

## Results

We investigated LinEs in both zygotic and azygotic meiosis but for different purposes. When incubated on sporulation medium, homothallic strains carrying the *h*^*90*^ mating-type allele switch mating-type, mate, and undergo zygotic meiosis (*i.e.*, meiosis immediately after mating) (Egel, 1977). Although such cultures are not synchronized for meiosis and thus not proper for phenotypic quantification, they reflect wild-type *S. pombe*, which is homothallic and undergoes zygotic meiosis. For phenotypic quantification, we used synchronized cultures of established diploids artificially induced for meiosis (*i.e.*, azygotic meiosis). We used two alleles, *pat1-114* (Iino and Yamamoto, 1985) and *pat1-as1* (Guerra-Moreno et al., 2012), to synchronize meiotic cultures by raising the temperature (to 34°C) or by adding an ATP analog (4-amino-1-tert-butyl-3-(3- methylbenzyl)pyrazolo[3,4- d]pyrimidine), respectively, to inactivate the Pat1 repressor of meiosis (Yamamoto, 1996). In this study, we used zygotic cultures (*h*^*90*^) for qualitative analyses and azygotic cultures (*pat1-114* or *pat1-as1*) for quantitative analyses.

### LinEs form long linear structures in zygotic meiosis in 25°C

We observed LinE morphology in live zygotic cultures (*h*^*90*^ strains) at 25°C using super-resolution microscopy (structured illumination microscopy, or SIM) and concentrated on cells in the horsetail stage (*i.e.*, meiotic prophase; Ding et al., 2004). We found that each LinE – Rec10-GFP, Rec25-GFP, Rec27-GFP and Mug20-GFP – showed long linear nuclear structures (Figures 1A and S1). These LinE forms were more nearly continuous than those in previous observations (Lorenz et al., 2004; Davis et al., 2008; Estreicher et al., 2012; Fowler et al., 2013). The morphology and length of the Rec25-GFP, Rec27-GFP and Mug20-GFP structures were very similar, while the morphology of Rec10-GFP was not as continuous as the others. Some of the horsetail nuclei showed more dotty LinE foci, suggesting that LinEs may assume different forms in different meiotic stages. Our observations are consistent with previous unpublished observations of the four LinE-GFP derivatives in *h*^*90*^ strains at 25°C (D-C Ding, personal communication) and indicate that the LinE complex may form end-to-end chromosomal structures like the SC.

**Figure 1.**
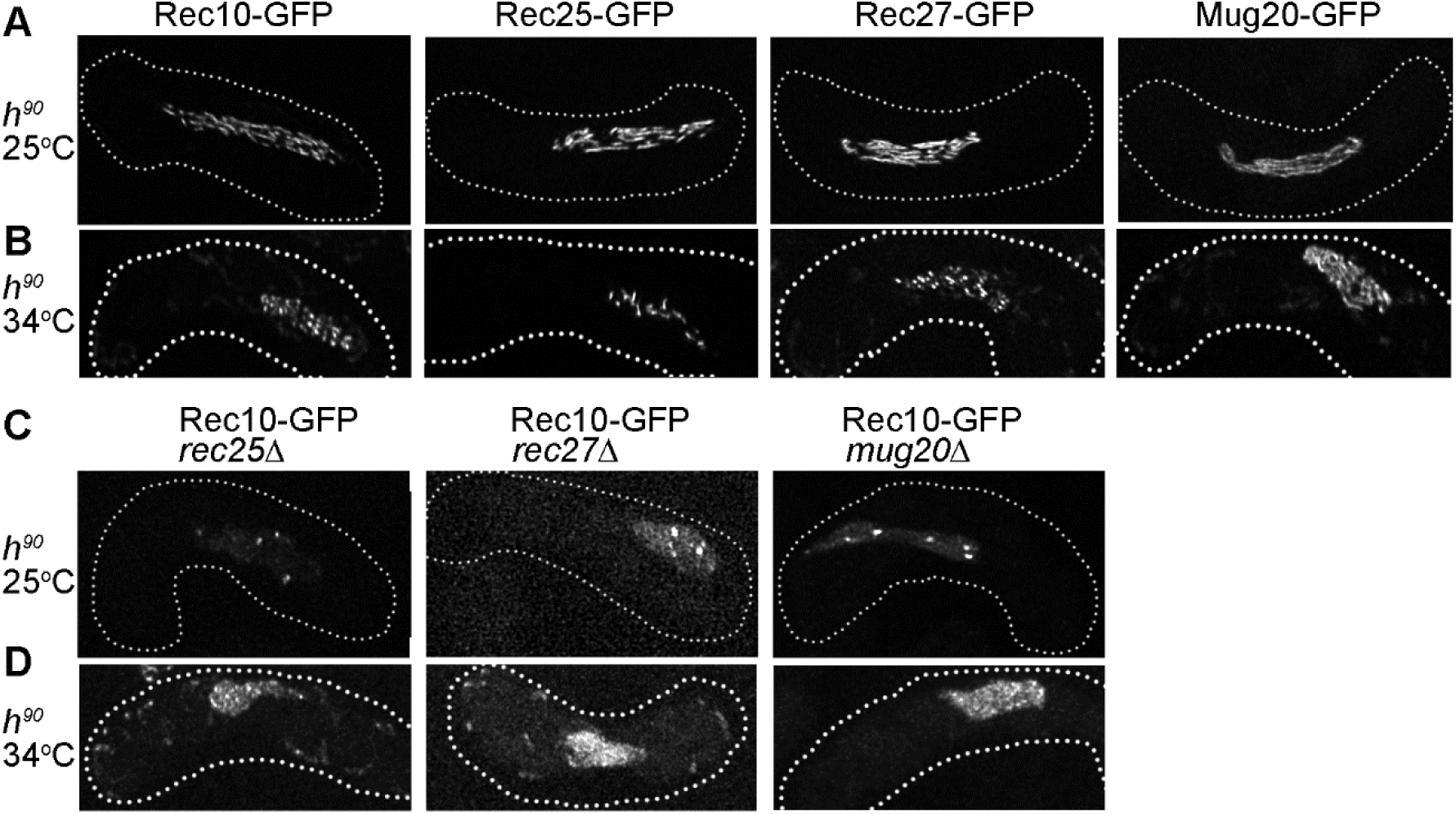
LinE subunit structures in live, zygotic meiosis. Rec10-GFP, Rec25-GFP, Rec27-GFP and Mug20-GFP were observed in *h*^*90*^ strains using Structured Illumination Microscopy (SIM). Each strain was incubated on sporulation medium (MEA) at 25°C for 13 – 15 hr or at 34°C for 10 – 17 hr. Each image is representative of at least 20 cells in the horsetail stage (*i.e*., meiotic prophase; Ding et al., 2004) for each strain. **(A)** LinE subunits form linear structures in zygotic meiosis at 25°C. All four LinE subunits formed continuous, linear structures. Each image shows the maximum intensity projection (maximal projection) of the entire Z-stack of image sections, shown in Figure S1. **(B)** Rec10-, Rec27- and Rec25-GFP fail to form linear structures at high temperature (34°C). Rec10-, Rec-25- and Rec27-GFP formed non-continuous structures and often showed bright nuclear dots. These “dotty foci” were different from the “linear structures” observed at 25°C (Figure 1A) but were similar to the structures in previous studies of azygotic meiosis at 30°C (Lorenz et al., 2004) or 34°C (Davis et al., 2008; Estreicher et al., 2012; Fowler et al., 2013). Mug20-GFP showed long linear structures at 34°C, similar to those at 25°C (Figure 1A). Each image shows the maximal projection of the entire Z-stack of image sections, shown in Figure S2. **(C)** In LinE mutants, Rec10-GFP enters the nucleus and forms a few bright nuclear foci and more nearly uniform nuclear distribution than in wt at 25°C in zygotic meiosis (Figure 1A). These results indicate that Rec10-GFP still enters the nucleus and forms nuclear foci, but not linear structures, in the absence of any other LinE subunit. Each image shows the maximal projection of the entire Z-stack of image sections, shown in Figure S4. **(D)** Rec10-GFP forms a few nuclear foci and nearly uniform nuclear distribution at 34°C in zygotic meiosis when other LinE subunits are missing. Each image shows the maximal projection of the entire Z-stack of image sections, shown in Figure S4. The dotted line represents the outline of the cell.

Previous published studies of LinE morphology were often done at 34°C using azygotic cultures (*pat1-114* strains) and in fixed cells (Davis et al., 2008; Estreicher et al., 2012; Fowler et al., 2013). These studies showed each LinE forms non-continuous, dotty foci. We next tested if the LinE linear forms are established at 25°C but not at 34°C in live *h*^*90*^ zygotic cells. In the 34°C cells in the horsetail stage, Rec10-GFP, Rec25-GFP and Rec27-GFP all showed dotty foci (Figures 1B and S2). These 34°C foci were dimmer than those at 25°C, and they were in non-continuous forms. Mug20-GFP foci still showed continuous linear forms at 34°C, but they were fuzzier than those at 25°C. The data indicate that LinEs form long linear structures in zygotic meiosis and that Rec10, Rec25 and Rec27 linear forms are sensitive to high temperature.

### Rec10-GFP forms nuclear foci independent of other LinE subunits

Previous studies using fluorescence microscopy of either fixed cells or nuclear spreads showed that nuclear focus-formation by each LinE depends on each of the other LinEs tested (Davis et al., 2008; Fowler et al., 2013). These studies were done in synchronized azygotic cultures (*pat1-114* strains) at 34°C, so we tested this genetic dependence in zygotic cultures (*h*^*90*^ strains) at 25°C. We found that Rec25-GFP, Rec27-GFP and Mug20-GFP did not form foci when any other LinE subunit was missing (Figure S3), showing their focus-formation is dependent on other LinE subunits, as previously reported under different conditions. However, Rec10-GFP formed a limited number of abnormal dotty nuclear foci in *rec25Δ, rec27Δ* or *mug20Δ* strains (Figures 1C and S4). The Rec10-GFP foci in these deletion strains were fuzzier and often showed a few bright dots in the nucleus; in addition, there was higher fluorescence spread nearly uniformly throughout the nucleus. This result shows that Rec10-GFP can form nuclear foci in *rec25Δ, rec27Δ* or *mug20Δ*, but the Rec10-GFP foci appear less continuous than those in WT.

These results differ from previous reports that Rec10-GFP does not form nuclear foci in *rec25Δ, rec27Δ* or *mug20Δ* (Davis et al., 2008; Fowler et al., 2013). The difference may result from the type of meiotic culture (azygotic *vs.* zygotic), the preparation of the sample (fixed *vs.* live cells) or the temperature (34°C *vs.* 25°C). Therefore, we examined Rec10-GFP foci in *rec25Δ, rec27Δ* or *mug20Δ* in *h*^*90*^ strains at 34°C. We found Rec10-GFP still formed a few nuclear foci in *rec25Δ, rec27Δ* or *mug20Δ* at 34°C (Figures 1D and S5). Our results indicate that in live, zygotic cultures Rec10 can bind to chromosomes and form limited abnormal dotty foci even without the other LinE subunits.

### Rec27-GFP forms long linear structures in azygotic meiosis at 25°C

LinEs may form long linear structures only in zygotic meiosis (*h*^*90*^ strains) but not in azygotic meiosis (*pat1-114* or *pat1-as1* strains). To test this hypothesis, we investigated the morphology of a representative LinE (Rec27-GFP) under different azygotic and temperature culture conditions (Figures 2 and S6). We found that Rec27-GFP formed dotty foci in *pat1-114* at 34°C (Figure S6A), which is consistent with previous studies (Davis et al., 2008; Fowler et al., 2013). Focus formation started at 2 hr after meiotic induction and ended around 4.5 hr, before MI (Figures 3A and 3B). Rec27-GFP also formed dotty foci in *pat1-as1* at 34°C (Figure S6B). The dotty foci in *pat1-as1* were slightly dimmer than those in *pat1-114*, but the focus number and formation timing were similar (Figure S6). These results suggest that the LinE focus-formation phenotype is similar in temperature-sensitive and analog-sensitive *pat1* mutants at 34°C, as expected from inactivation of the same regulatory protein (Yamamoto, 1996).

**Figure 2.**
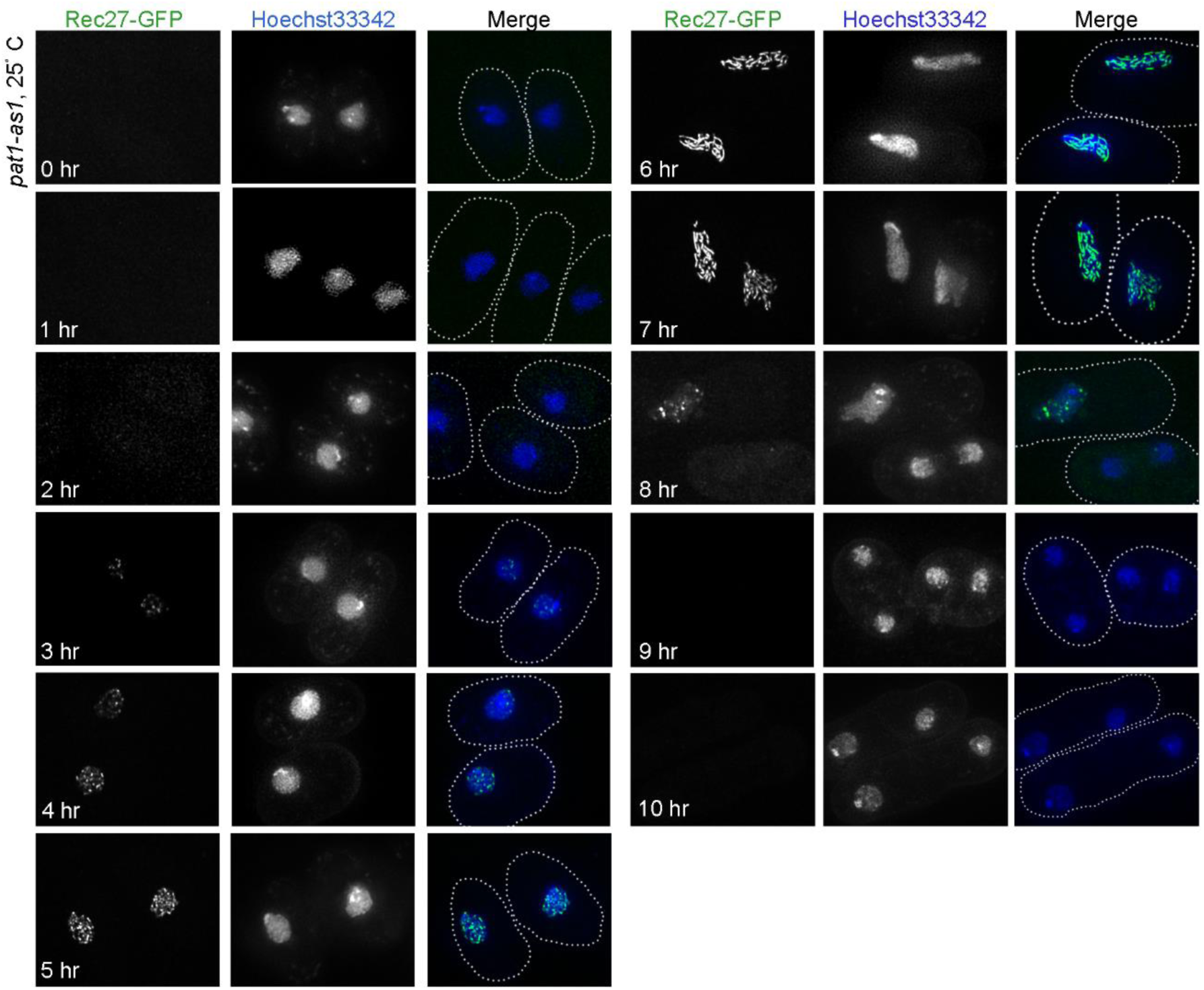
Rec27-GFP changes conformation during synchronized azygotic meiosis at 25°C or 34°C. Rec27-GFP was observed in live cells of synchronized, azygotic meiotic cultures (*pat1-as1*) at 25°C. Cells were collected at the indicated times after meiotic induction and were observed with Structured Illumination Microscopy (SIM). Each image is representative of at least 50 cells analyzed in each condition. Rec27-GFP “dotty” foci appeared at 3 hr after meiotic induction and increased until 7 hr when they became “linear structures”, showing a conformational transition of Rec27-GFP. The linear structures at 7 hr were shorter than those observed in zygotic meiosis at 25°C (Figure1A) but were much more continuous than those observed in 34°C (Figures 1B and S6) (Lorenz et al., 2004; Davis et al., 2008; Estreicher et al., 2012; Fowler et al., 2013). Nuclear location is shown by Hoechst 33342 staining. The dotted line represents the outline of the cell.

**Figure 3.**
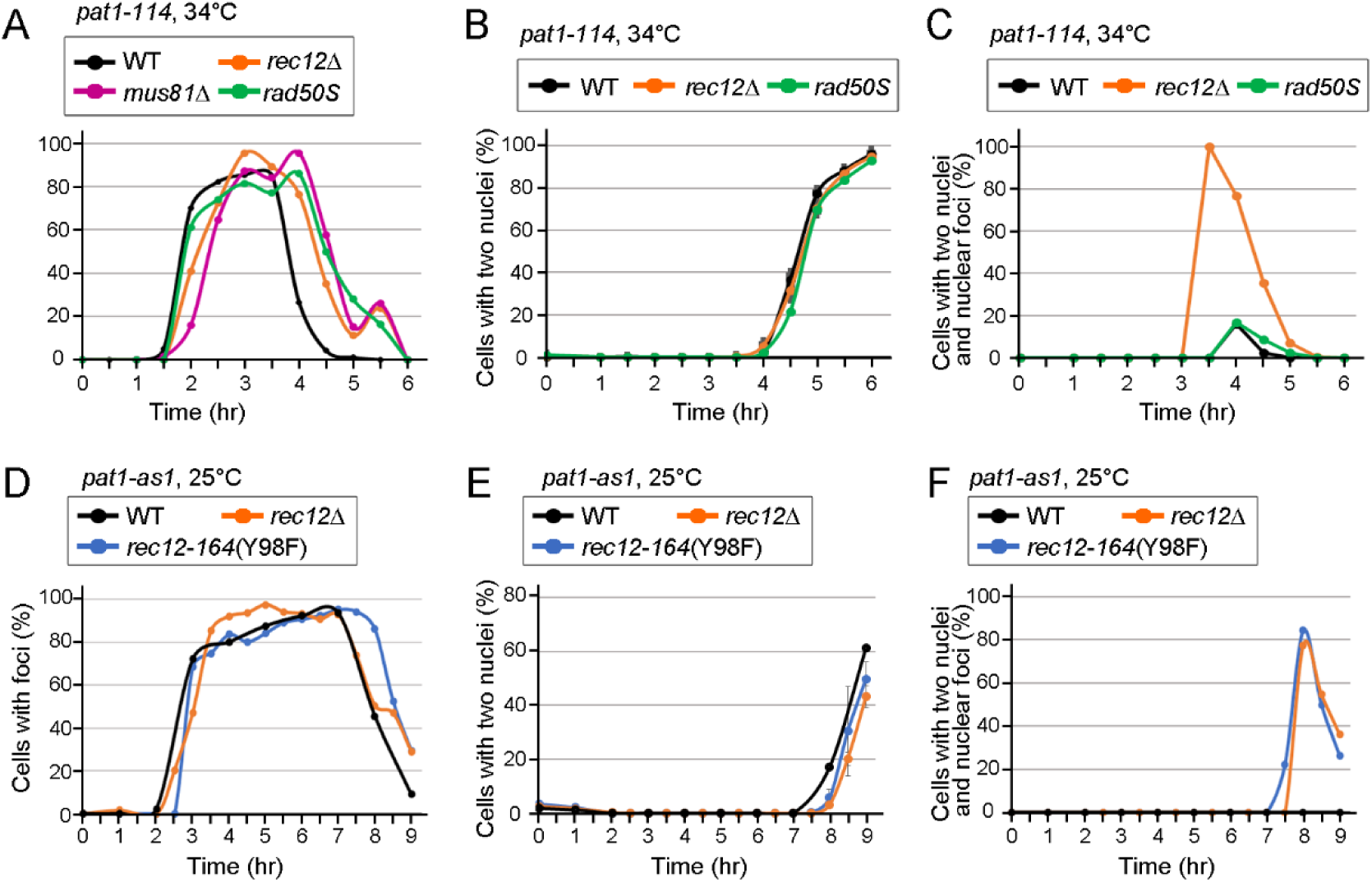
Rec27-GFP focus formation is not affected by DSB formation or repair, but DSB absence delays Rec27-GFP focus disassembly. Quantification of the Rec27-GFP focus analysis in various mutants in *pat1-114* (34°C) or *pat1-as1* (25° C) azygotic cultures. For each condition, over 250 randomly selected cells, in at least ten randomly selected microscopic fields, were examined. **(A-C)** Rec27-GFP focus analysis in *pat1-114* (34°C). **(A)** Rec27-GFP “dotty” foci appeared ∼3 hr after meiotic induction in WT, *rec12Δ, rad50s* and *mus81Δ*. The foci went away between 4 and 5 hr. **(B)** Quantification of cells with two or more nuclei in WT, *rec12Δ* and *rad50s*. **(C)** Quantification of cells with two nuclei but still having Rec27-GFP foci in one or both nuclei in WT, *rec12Δ* and *rad50s*. The numbers of observed cells with two nuclei in *rec12Δ* at 3.5 hr, 4 hr, 4.5 hr and 5 hr were 2, 33, 67 and 26, respectively. **(D-F)** Rec27-GFP focus analysis in *pat1-as1* (25°C). **(D)** Rec27-GFP structures appeared around 3 hr after meiotic induction and went away between 7 and 8 hr in WT, *rec12Δ* and *rec12-164* (Y98F). The timing of Rec27-GFP structure formation and disassembly was similar in WT and mutants. **(E)** Quantification of cells with two or more nuclei in WT, *rec12Δ* and *rec12-164* (Y98F). **(F)** Cells with delayed Rec27-GFP focus disassembly. Quantification of cells with two nuclei but still having Rec27-GFP structures in one or both nuclei in WT, *rec12Δ* and *rec12-164* (Y98F). The numbers of observed cells with two nuclei in *rec12Δ* at 8 hr, 8.5 hr and 9 hr were 15, 73, and 82, respectively. The number of observed cells with two nuclei in *rec12-164*(Y98F) at 8 hr, 8.5 hr and 9 hr were 43, 68, and 78, respectively. Both *rec12Δ* and *rec12-164*(Y98F) showed delay in Rec27-GFP structure disassembly.

We then investigated Rec27-GFP focus morphology in azygotic (*pat1-as1*) strains at 25°C (Figure 2). We found about half of the population showed a few Rec27-GFP foci at 3 hr after meiotic induction, before replication started (Figures 2, 3D and S7). Over 85% of the cells had Rec27-GFP foci at and after 5 hr, and the foci disassembled around 8 – 9 hr, before MI (Figures 2 and 3D). This result shows that Rec27-GFP foci exist for ∼5 hr at 25°C, likely due to the slower cell cycle progression at 25°C than at 34°C. In addition, most of the cells showed dotty foci at 5 hr, and the dotty foci turned into linear structures at 6 – 7 hr (Figure 2). The linear structures then became dotty again at about 8 – 9 hr and went away before MI at about 9 hr (Figure 2). This result indicates that LinEs form dot-like foci first and later expand along the chromosome into a linear form, before returning to the dot-like form. In addition, linear LinE forms were found only at 25°C, not at 34°C, indicating that formation of the linear-expansion structure is temperature-sensitive.

### LinE foci are sensitive to 1,6-hexanediol, like the SC in budding yeast, flies, and worms

The linear form of LinEs suggested that *S. pombe* forms SC-like structures in zygotic meiosis at 25°C when examined in live cells. To test this possibility, we tested if LinEs share a notable character of the SC. The SC in budding yeast, flies, and worms shows liquid-crystalline properties, as indicated by sensitivity of the SC, but not cohesin, to the chaotropic agent 1,6-hexanediol (Rog et al., 2017). We found that the LinE linear forms were not visible after treating the cells with 10% 1,6-hexanediol for five min (Figures 4 and S8). All four LinE subunits, separately tagged with GFP, showed similar sensitivity to the treatment. Over 90% of treated cells did not show the LinE linear structure; they showed nearly uniform nuclear signal after 1,6-hexanediol treatment, while some of them showed a few dots in the nucleus. We quantified nuclear GFP intensity of all four LinE-GFPs, before and after treatment. Nuclear Rec10-GFP, Rec25-GFP and Rec27-GFP intensity was reduced to 50 – 60% after the treatment, while Mug20-GFP was reduced to 30% (Figure S9). Some treated cells still showed high nuclear GFP intensity but none of them showed LinE linear structures (Figures 4 and S8). In all four LinE-GFP cells, Rec8, a meiotic cohesin subunit, remained the same in those treated cells, indicating that LinEs but not cohesins are sensitive to 1,6-hexanediol treatment (Figure 4).

**Figure 4.**
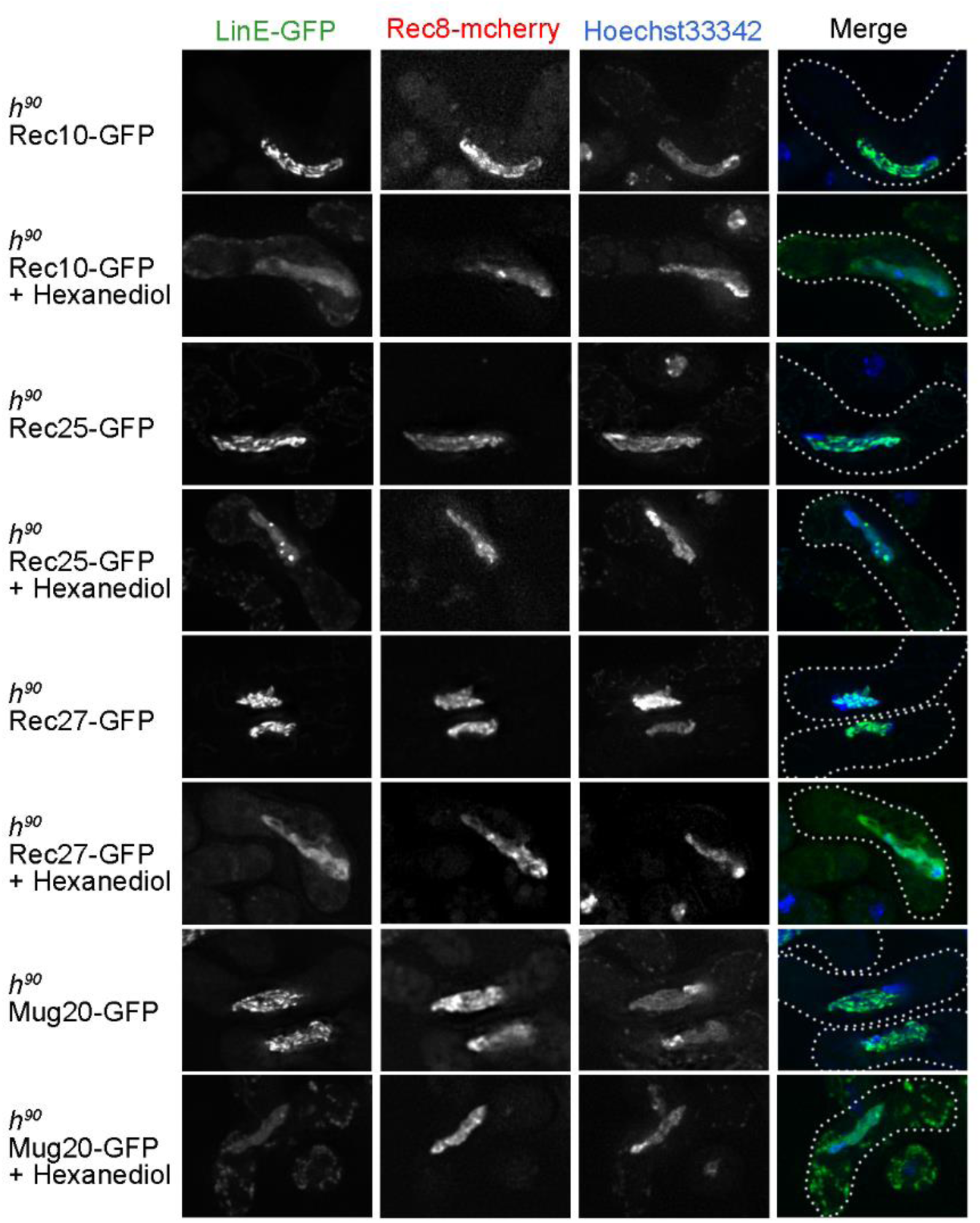
LinE subunits but not cohesin subunit Rec8 are sensitive to 1,6-hexanediol treatment. Rec10-GFP, Rec25-GFP, Rec27-GFP and Mug20-GFP were observed in *h*^*90*^ strains with and without 1,6-hexanediol treatment. Cells were incubated at 25°C for 14 – 16 hr and observed in high-resolution fluorescence microscopy (DeltaVision). 1,6-Hexanediol (10% in EMM2-N) was added for five min at room temperature before observation. After treatment, Rec10-GFP, Rec25-GFP, Rec27-GFP and Mug20-GFP lost their nuclear localization or became lumpy, but cohesin subunit Rec8-mCherry was little changed. These results show that LinE structure is sensitive to 1,6-hexanediol, a character of the SC (Rog et al., 2017). Nuclear localization is shown by Hoechst 33342 staining. The merge column shows the merged image of LinE-GFP and Hoechst 33342 staining. The dotted line represents the outline of the cell.

### LinE focus-formation is independent of meiotic DSB formation and DSB repair, but lack of meiotic DSB formation delays LinE focus disassembly

Previous studies using nuclear spreads showed that Rec10-GFP forms foci in *rec12Δ* (Lorenz et al., 2004; Davis et al., 2008), indicating that meiotic DSB formation is not needed for LinE focus-formation. To understand which meiotic event(s) is (are) necessary for LinE focus-formation and disassembly, we analyzed Rec27-GFP foci in synchronized azygotic cultures in three meiosis-deficient mutants: *rec12Δ* (DSB formation-deficient (Cervantes et al., 2000), *rad50S* (DSB resection-deficient; Young et al., 2002), and *mus81Δ* (Holliday junction resolution-deficient; Boddy et al., 2001; Smith et al., 2003; Cromie et al., 2007) in azygotic *pat1-114* at 34°C. The Rec27-GFP focus-formation timing, focus number, and focus morphology were similar in these mutants and the wild-type control (Figures 3A and S6), showing that DSB formation and repair are not essential for LinE focus formation.

We also compared the Rec27-GFP focus disassembly in mutants and wild-type control in azygotic *pat1-114* at 34°C and found that failure of DSB formation leads to a LinE focus-disassembly defect (Figure 3C). In *rec12Δ* (*pat1-114* at 34°C), Rec27-GFP foci formed at the same time as in wild type, but over 50% of the *rec12Δ* cells with two nuclei showed foci in one or both nuclei at 3.5 hr and 4 hr (Figure 3C). However, no Rec27-GFP foci were found in cells with four nuclei (which began at 5 hr) (Figure 3C and data not shown), showing that Rec27-GFP focus disassembly was delayed in some cells until after MI but happened before the second meiotic division (MII) in *rec12Δ*. This phenotype was also found in *rec12Δ pat1-as1* at 25°C (Figure 3D-F), suggesting the focus-disassembly defect is not related to temperature or meiotic induction method differences. To test if the delayed disassembly of LinE foci is related to DSB formation rather than some other feature of Rec12, we did the same analysis in *rec12-164* (Y98F), an active-site mutation (Cervantes et al., 2000). *rec12-164* (Y98F) showed the same delayed disassembly of Rec27-GFP foci as *rec12Δ* (Figure 3D-F). These results showed that meiotic DSB formation is necessary for timely LinE focus disassembly but not for their assembly.

We also investigated the Rec27-GFP disassembly in DSB repair-deficient mutants. Rec27-GFP foci disassembled normally in the *rad50S* mutant, which is deficient in removal of Rec12 covalently bound to DSBs that it forms (Cromie et al., 2007) (Figure 3A). Rec27-GFP focus disassembly was delayed in the *mus81Δ* HJ resolution-deficient mutant (data not shown), but the cells were arrested before MI entry (Boddy et al., 2001).

### LinE missense mutants have defective LinE structures

Several studies have shown how LinE subunits interact with each other and other meiotic proteins using yeast two-hybrid analysis or immunoprecipitation analysis (Loidl, 2006; Spirek et al., 2010; Miyoshi et al., 2012; Estreicher et al., 2012; Sakuno and Watanabe, 2015). We and others have shown that all LinE subunits are needed for intact LinE complex formation (Figures 1C and S3) (Davis et al., 2008; Fowler et al., 2013). However, how the LinE subunits interact and how the LinE complex is formed are not well understood. To address this question, we used *h*^*90*^ strains at 25°C to assay LinE focus-formation in LinE subunit missense mutants that show defects in recombination and meiotic DSB formation (Ma et al., 2017) (Figures 5 and S10). Based on their LinE focus phenotype, we classified the LinE mutants into three groups. **a)** *rec27-238* (K28E) showed dot-like Rec25 foci. Thus, in *rec27-238* (K28E) the LinE (Rec25-GFP) could enter the nucleus and interact with other LinEs but did not form intact, linear structures. **b)** *rec25-236* (K124I), *rec27-239* (R32E), *rec27-240* (K36E) and *mug20-251* (V78E) showed much more uniform Rec25-GFP fluorescence throughout the nucleus. There were a few nuclear GFP foci in these mutants, but fewer than in group **a**. Thus, in these mutants the LinEs could enter the nucleus but seemed not to interact with other LinEs to form abundant foci or linear structures. **c)** *mug20-250* (L52P) showed very few nuclear Rec25-GFP signals, suggesting this mutant protein is defective in Rec25 binding or in LinE complex formation. All of these LinE mutants show a defect in meiotic DSB formation and recombination (Ma et al., 2017). Moreover, group **b** mutants show more severe defects than group **a** mutants, and DSBs are undetectable in group **c** mutant *mug20-250* (Ma et al., 2017). We also assayed Rec25-GFP formation in *rec27-238* and *rec27-240* in azygotic meiosis (*pat1-as1*) by the same method used in Figure 2; Rec25-GFP never formed linear structures through the whole time-course (Figure S11). These data indicate a strong correlation between DSB formation, meiotic recombination, and LinE linear structure establishment.

**Figure 5.**
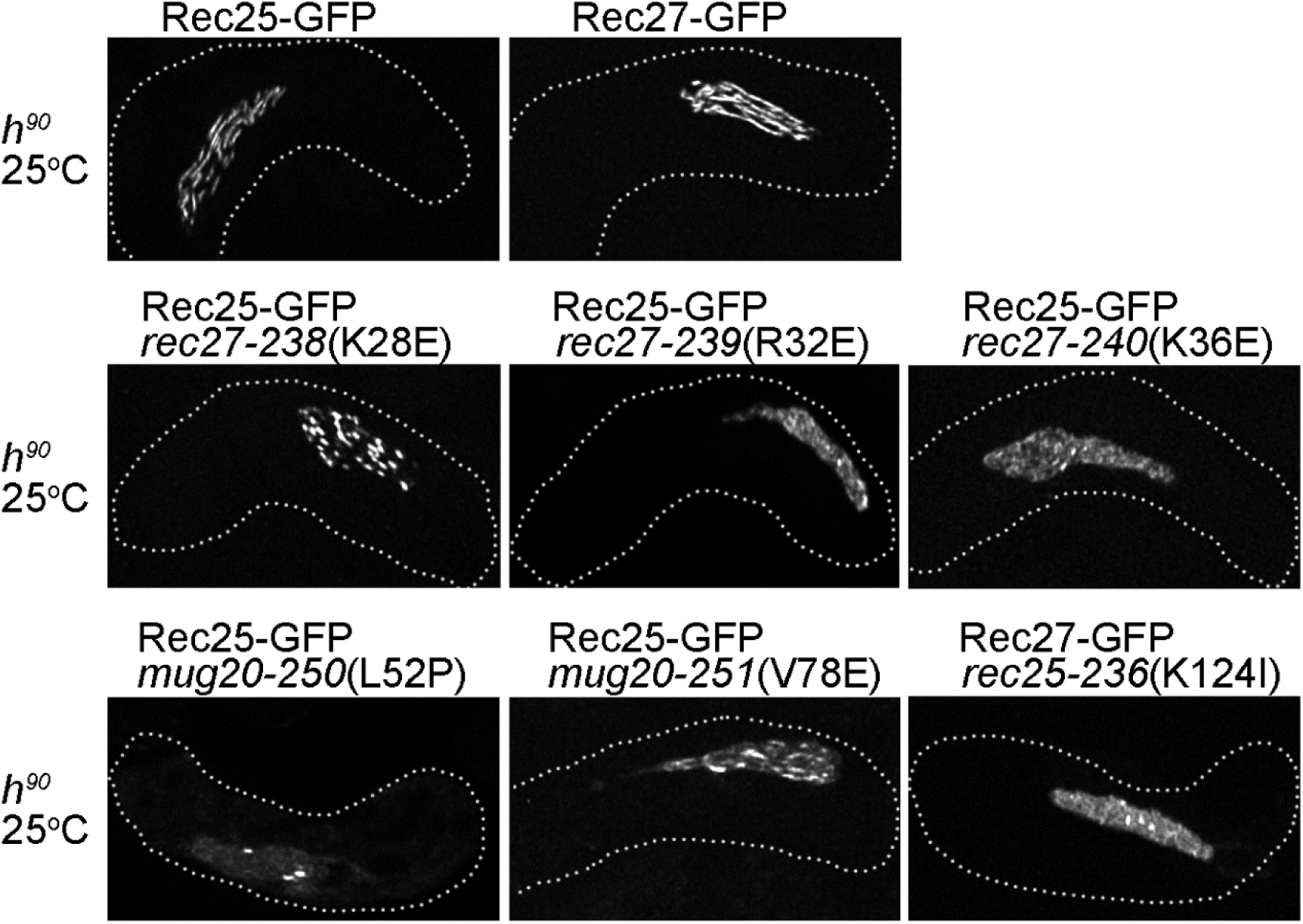
LinE missense mutations alter LinE linear structure. Rec25-GFP (in WT, *rec27* and *mug20* missense mutants) and Rec27-GFP (in a *rec25* missense mutant) were analyzed in live zygotic cultures at 25°C. Cells were incubated on MEA and at least 20 cells for each strain were analyzed using high-resolution fluorescence microscopy (DeltaVision). None of the mutants formed intact LinE linear structures. In *rec27-238, rec27-239* and *mug20-251*, Rec25-GFP showed non-continuous dotty foci. Rec25-GFP in *rec27-240* and Rec27-GFP in *rec25-236* showed a few dot-like foci, but Rec25- or Rec27-GFP still entered the nucleus. Rec25-GFP in *mug20-250* formed very few, if any, dot-like foci. Nuclear localization is shown by Hoechst 33342 staining. The dotted line represents the outline of the cell.

## Discussion

Previous studies have concluded that *S. pombe* does not form an SC in meiosis (Olson et al., 1978; Dresser and Giroux, 1988; Bähler et al., 1993). LinE subunits Rec10, Rec25, Rec27 and Mug20 have been reported to form dot-like or short linear structures in meiosis, not end-to-end chromosomal structures resembling the SC of all other species examined (except for *Aspergillus nidulans*) (Egel-Mitani et al., 1982; Bähler et al., 1993; Lorenz et al., 2004; Davis et al., 2008; Estreicher et al., 2012; Fowler et al., 2013). However, in most of these studies, cells were from azygotic cultures at high temperature (34°C) and were chemically fixed. Studies of LinE morphology with live cells from zygotic cultures at the physiologically optimal temperature (about 25°C) are limited (Fowler et al., 2013). In this study, we analyzed LinE foci in live cells in zygotic meiotic cultures at 25°C and azygotic meiotic cultures at both 25°C and 34°C. We found that LinEs form end-to-end chromosomal structures in wild-type (*h*^*90*^) cultures at 25°C, suggesting that *S. pombe* forms structures that may be equivalent to the SC of other species (Figure 1A) (see Introduction). We propose that the SC in *S. pombe* is more fragile and collapses at high temperature, or upon fixation or nuclear spreading. In addition, LinE structures are sensitive to 1,6-hexanediol treatment just like the SC in *Saccharomyces cerevisiae, Drosophila. melanogaster* and *Caenorhabditis elegans* (Rog et al., 2017) (Figure 4). This result also supports *S. pombe* forming SC or SC-like structures in meiosis.

Our data showed that the morphologies of Rec25-, Rec27- and Mug20-GFP foci in zygotic culture at 25°C were quite similar, while Rec10-GFP foci showed several differences (Figures 1A, 1C, S1 and S4). All four LinE subunits formed linear structures (Figures 1A and S1), but the Rec10-GFP linear structures were slightly shorter than the others. In addition, previous studies showed the DNA binding preference of Rec10 is different from those of the other LinE subunits: Rec25, Rec27 and Mug20 are concentrated at DSB hotspots, whereas Rec10 is more uniformly distributed along chromosomes (Miyoshi et al., 2012; Fowler et al., 2013; Kariyazono et al., 2019). Our observation of a Rec10-GFP morphology difference may reflect the Rec10 distribution. Moreover, in the absence of other LinEs we found Rec10-GFP showed a more uniform nuclear distribution with a few nuclear foci (Figures 1C and S4). This result supports the idea that Rec25, Rec27 and Mug20 enhance Rec10’s binding to DSB hotspots. We also found that Rec10-GFP entered the nucleus and formed nuclear foci in other LinE subunit deletion mutants at either 25°C or 34°C (Figures 1C-D, S4 and S5). This result is not consistent with previous studies, which showed that Rec10-GFP does not form visible nuclear foci in *rec25Δ, rec27Δ* or *mug20Δ*; these studies were done with fixed, azygotic cells (Davis et al., 2008; Fowler et al., 2013). However, we observed that Rec25-, Rec27- and Mug20-GFP do not detectably appear in the nucleus in any of the LinE subunit deletion mutants (Figure S3). This result suggests that, unlike Rec10-GFP, the localizations of Rec25, Rec27 and Mug20 are interdependent with the other LinE subunits. These results support the model that the four LinE subunits form a complex in the cytoplasm and that this complex is transported into the nucleus by the nuclear localization signal on Rec10 (Lorenz et al., 2004; Wintrebert et al. 2020).

We found two configurations of LinEs: dotty foci and linear structures. The dotty foci appeared before the time of replication (Figure S7), then became linear around the time of DSB formation (Hyppa et al., 2014), and finally returned to dotty foci before disassembling just before MI (Figures 2, 3 and S7). Previous studies showed that with faster kinetics at 34°C LinEs form dotty foci 3.5 – 4.5 hr after meiotic induction (Davis et al., 2008; Fowler et al., 2013), indicating that the dotty LinEs, seen by microscopy, may be closely related to the LinE clusters inferred from 3C analyses of cells induced for meiosis under the same conditions (Fowler et al., 2018). We propose that LinE clusters also undergo a conformational change during meiotic progression. LinEs gather nearby clusters into a visible dotty focus, which may expand to fill the chromosomal gaps between the foci and become the linear structure. In addition, LinEs likely remain on the chromosome after the DSBs are made and until MI (Figures 2, 3D, and S7; Hyppa et al., 2014). This proposal raises the possibility that LinEs also participate in other meiotic processes, such as DSB repair, including chromosomal partner choice for DSB repair. Linear structures were not found at 34°C (Figures 1B and S2), suggesting that either the formation of linear structures is temperature-sensitive or they are too transient for us to observe at 34° C. Previous studies of genome-wide DSBs showed similar DSB profiles at both 34°C and 25°C (Hyppa et al., 2014); however, several DSB hotspots were found only at 34°C or at 25°C, indicating temperature may change the LinE binding pattern at certain loci. Determining the difference between the two LinE configurations and understanding the regulatory mechanism of the configuration change will likely shed light on the role of LinEs in DSB interference and DSB repair.

LinE focus disassembly was delayed in both *rec12Δ* and *rec12-164* (Y98F), showing that DSB formation is needed for a later event, proper LinE disassembly. However, those LinE foci remaining in cells with two nuclei went away before MII, which begins at 5 hr at 34°C or 8 – 9 hr at 25°C (Figures 3A and 3D). This result indicates that meiotic DSB formation is one of the factors leading to LinE focus disassembly, but there are other mechanisms regulating the process. Since LinEs determine meiotic DSB hotspots and are necessary for DSB formation at most hotspots (Fowler et al., 2013), we propose that LinEs stay on the chromosomes until they sense that sufficient DSBs have been formed. But another mechanism can remove LinE clusters before MII, allowing the sister chromatids to complete their separation at MII. Several proteins, such as Rec15 and Hop1, interact with both Rec10 and the DSB forming complex (Kariyazono et al., 2019) and may also be involved in LinE structure disassembly.

Based on our observations, we propose a model of the two sequential dynamic configurations of LinEs (Figure 6). Rec10, Rec25, Rec27 and Mug20 form a LinE complex in the cytoplasm and are transported into the nucleus (Figure 6A). After DNA replication, LinEs bind to meiotic DSB hotspots, form DSB hotspot clusters, which appear as dotty foci in light microscopy of live cells, and promote DSB formation (Figure 6B). LinEs than expand along the chromosome, making linear structures that may participate in other meiotic processes, such as some aspect of DSB repair. Before MI, the LinE linear structures disassemble and show dotty foci again (Figure 2; 8 hr). Finally, the LinE complex dissociates from the chromosomes. Missense mutants of two LinE components, Rec25 and Rec27, showed only the first configuration (dotty foci) (Figures 5, S10, and S11) but failed to undergo the dotty-to-linear transition. This result indicates these point mutations interrupt a critical interaction between LinE complexes. The recombination-deficient phenotype of these mutants implies that the linear form of the LinE complex is important for proper meiosis. Our results reveal two configurations of LinE structures, “dotty” foci and “linear structures”, which may represent two faces of LinEs in regulating meiotic recombination.

**Figure 6.**
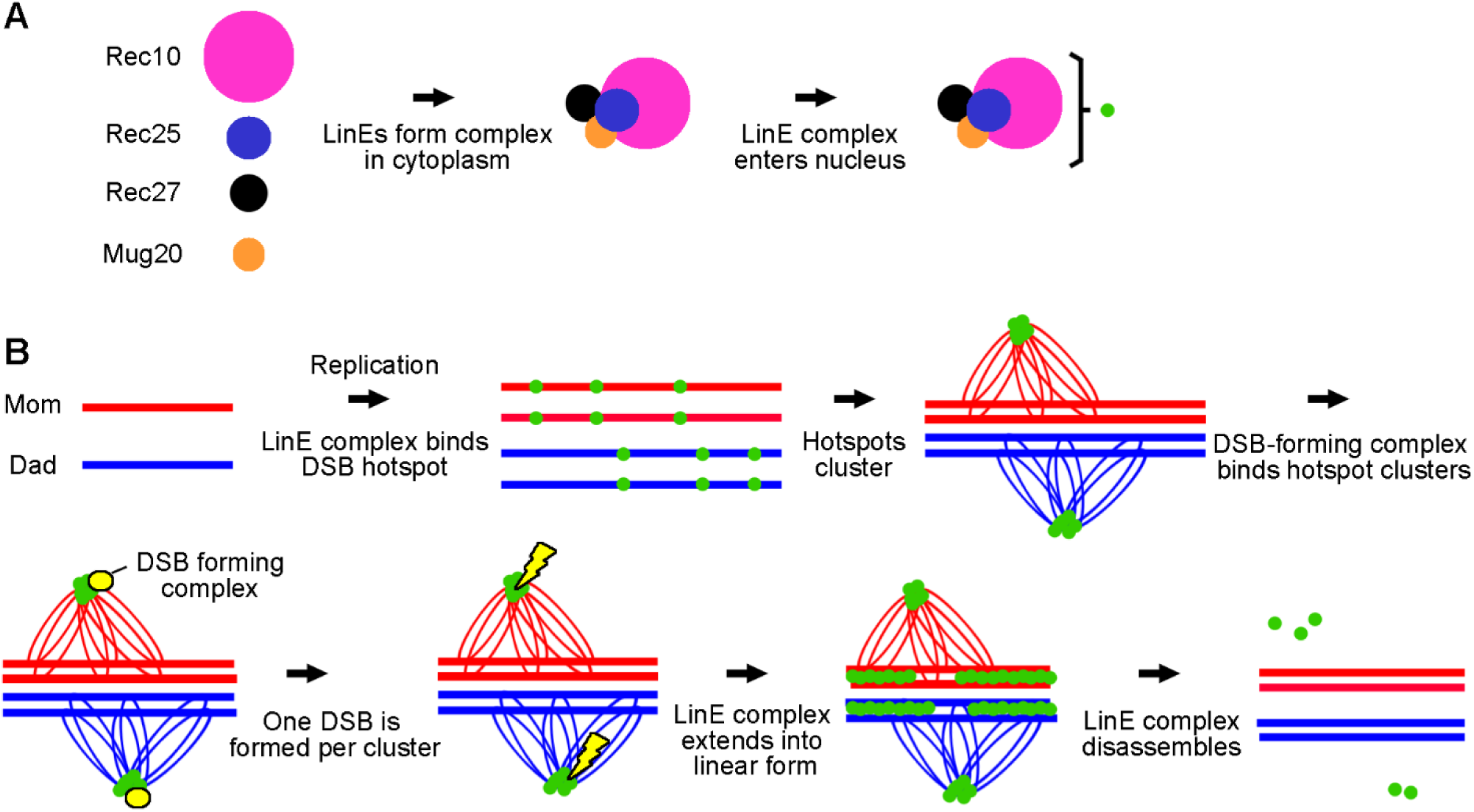
Model of two configuations of LinEs. **(A)** In the cytoplasm Rec10, Rec25, Rec27 and Mug20 form the LinE complex, which is transported into the nucleus via the nuclear localization signal on Rec10. **(B)** The transition of two LinE configurations. LinE complexes bind to DSB hotspots after DNA replication, and LinE-bound hotspots within an ∼200 kb region form a cluster (Fowler et al., 2018); a few clusters assemble into a “dotty” focus visible in the microscope. The DSB-forming complex is activated by the LinEs in a cluster and introduces one meiotic DSB in each DSB hotspot cluster. After DSB formation, the LinE complex extends along the chromosome, forming a “linear structure” visible in the microcope. The linear structures may aid DSB repair, perhaps by regulating partner choice for DSB repair. The LinE complex dissociates from the chromosomes before MI.

## Materials and Methods

### *S. pombe* strains and meiotic induction methods

*S. pombe* strains used in this study are listed in Supplementary Table S1. Media for cell growth and meiosis were as described (Smith, 2009). For zygotic meiotic induction (*h*^*90*^ strains), 50 – 100 μl of fresh overnight culture in YEL were collected and washed three times with water. Cells were suspended in water, spotted on sporulation medium (MEA) and incubated at 25°C for 13 – 15 hr or 34°C for 10 – 17 hr. After incubation, cells were collected and suspended in EMM2-N (water for Fig. 3), for microscopic assay. For azygotic meiotic induction (*pat1-114* or *pat1-as1* strains), cells were treated as described (Hyppa and Smith, 2009; Guerra-Moreno et al., 2012). At each meiotic time point, 500 μl of culture was collected for microscopic assay.

### Fluorescence microscopy

Cells induced for meiosis were suspended in EMM2-N with Hoechst 33342 (5 μg/ml). The cell suspension was spread on a poly-L-lysin-coated slide before observing by Structured Illumination Microscopy (SIM) with 100X/1.4 NA STED White Objective (Leica Microsystems) and VT-iSIM system (Visitech International), or DeltaVision Elite microscope with 100x/1.4 UPlanSApo lens (GE Healthcare). Image data were processed with Micro Evolution Deconvolution Software running in FIJI (SIM) or SoftWoRX (DeltaVision). Images are maximum intensity projections of sections with step size of 0.2 μm, to cover the whole cell. For cells treated with 1,6-hexanediol, cells were suspended in EMM2-N with Hoechst 33342 (5 μg/ml) and 1,6-hexanediol (10% w/v) and observed in a microscope within five minutes after addition of hexanediol. Experiments using two different sources of 1,6-hexanediol (Sigma-Aldrich and Hampton Research) showed similar results.

For quantification of LinE structure formation and disassembly, a cell suspension at each time point was spread on a poly-L-lysine-coated slide before observing in an EVOS FL Auto 2 inverted microscope (Thermo Fisher Scientific) with a 60X/1.42 PlanApo N Lens. Results using EMM2-N or water for suspending cells were not significantly different (data not shown). For each time point of each sample, at least 250 cells with clear nuclear staining from at least ten individual fields were examined. Three biological repeats were performed for each condition.

### Flow cytometry analysis

DNA content of azygotic meiosis samples were analyzed as described by Cervantes et al. (2000). One ml of meiotic culture was collected at each time point and fixed with 70% ethanol. Samples were treated by RNAase for 1-2 hr, suspended in 50 mM sodium citrate with 4 μg/ml propidium iodide and sonicated for 30 sec before analysis in FACSCanto™ II (Becton Dickinson).

### Quantification of nuclear LinE-GFP intensity

Cells with Rec10-GFP, Rec25-GFP, Rec27-GFP, or Mug20-GFP induced for zygotic meiosis (*h*^*90*^ strains) were examined as described above using DeltaVision Elite microscope, and sum slices of each image were projected with step size of 0.2 μm, to cover the whole cell. Nuclear region was identified by Hoechst 33342 staining. Nuclear GFP was quantified using ImageJ (NIH), and the nuclear region threshold was adjusted by Li method (Li and Lee, 1993; Li and Tam, 1998; Sezgin and Sankur, 2004).

## Acknowledgements

We are especially grateful to Da-Qiao Ding for sending us unpublished micrographs of live *S. pombe* zygotic meiotic cells expressing LinE-GFP proteins in the linear configuration; Yoshinori Watanabe for a strain with the *rec8*^*+*^*-mCherry-Tspo5<<nat*^*r*^ allele; Kami Ahmad for use of his EVOS FL Auto 2 inverted microscope; and Sue Amundsen, Randy Hyppa, Mai-Chi Nguyen, and especially Cristina Martín-Castellanos for helpful comments on the manuscript. This research was supported by grants R35 GM118120 to G.R.S. and P30 CA015704 to the Fred Hutchinson Cancer Research Center from the National Institutes of Health of the United States of America.

