## Supplemental information for "Dynamic configurations of meiotic hotspot determinants"

### Supplementary Table S1. *S. pombe* strains

| Strain | Genotype[a] | Used in |
|---|---|---|
| GP6966 | *$h^-/h^-$ ade6-3049/ade6-3049 pat1-114/pat1-114 rec27-205-GFP::kanMX6/rec27-205-GFP::kanMX6 lys4-95/+ +/his4-239* | Figure 3A-C, S6A |
| GP8762 | *$h^{90}$ rec10-203-GFP::kanMX6* | Figure 1A-B, S1-3 |
| GP8766 | *$h^{90}$ rec25-204-GFP::kanMX6* | Figure 1A-B, 5, S1-3 |
| GP8819 | *$h^{90}$ rec27-205-GFP::kanMX6* | Figure 1A-B, 5, S1-3 |
| GP8829 | *$h^{90}$ mug20-GFP::kanMX6* | Figure 1A-B. S1-3 |
| GP9506 | *$h^{90}$ rec25-204-GFP::kanMX6 rec27-238* | Figure 5, S11 |
| GP9508 | *$h^{90}$ rec25-204-GFP::kanMX6 rec27-240* | Figure 5, S11 |
| GP9529 | *$h^{90}$ rec25-204-GFP::kanMX6 rec27-239* | Figure 5 |
| GP9549 | *$h^{90}$ rec25-204-GFP::kanMX6 rec27-184::kanMX6* | Figure S3 |
| GP9553 | *$h^{90}$ rec25-204-GFP::kanMX6 mug20::natMX6* | Figure 5 |
| GP9554 | *$h^{90}$ rec25-204-GFP::kanMX6 mug20-251* | Figure 5 |
| GP9581 | *$h^{90}$ rec25-204-GFP::kanMX6 mug20-250* | Figure 5 |
| GP9591 | *$h^{90}$ rec27-205-GFP::kanMX6 rec25-180::kanMX6* | Figure S3 |
| GP9592 | *$h^{90}$ rec10-203-GFP::kanMX6 rec25-180::kanMX6* | Figure 1C-D, S4-5 |
| GP9593 | *$h^{90}$ mug20-GFP::kanMX6 rec25-180::kanMX6* | Figure S3 |
| GP9594 | *$h^{90}$ rec27-205-GFP:: kanMX6 mug20::natMX6* | Figure S3 |
| GP9595 | *$h^{90}$ mug20-GFP::kanMX6 rec27-184::kanMX6* | Figure S3 |
| GP9596 | *$h^{90}$ rec10-203-GFP::kanMX6 rec27-184::kanMX6* | Figure 1C-D, S4-5 |
| GP9597 | *$h^{90}$ rec27-205-GFP::kanMX6 rec10-175::kanMX6* | Figure S3 |
| GP9598 | *$h^{90}$ rec25-204-GFP::kanMX6 rec10-175::kanMX6* | Figure S3 |
| GP9599 | *$h^{90}$ mug20-GFP::kanMX6 rec10-175::kanMX6* | Figure S3 |
| GP9600 | *$h^{90}$ rec10-203-GFP:: kanMX6 mug20::natMX6* | Figure 1C-D, S4-5 |
| GP9603 | *$h^-/h^-$ ade6-3049/ade6-3049 pat1-114/pat1-114 rec27-205-GFP::kanMX6/rec27-205-GFP::kanMX6 rad50S/red50S lys4-95/+ +/his4-239* | Figure 3A-C |
| GP9604 | *$h^-/h^-$ ade6-3049/ade6-3049 pat1-114/pat1-114 rec27-205-GFP::kanMX6/rec27-205-GFP::kanMX6 mus81::kanMX/mus81::kanMX lys4-95/+ +/his4-239* | Figure 3A |
| GP9606 | *$h^-/h^-$ ade6-3049/ade6-3049 pat1-114/pat1-114 rec27-205-GFP::kanMX6/rec27-205-GFP::kanMX6 tel1::kanMX/tel1::kanMX rec12-169::kanMX/rec12-169::kanMX lys4-95/+ +/his4-239* | Figure 3A |
| GP9679 | *$h^{90}$ rec27-205-GFP::kanMX6 $rec8^+$-mCherry-$T_{spo5}$<<$nat^r$* | Figure 4, S8-9 |
| GP9702 | *$h^{90}$ rec10-203-GFP::kanMX6 $rec8^+$-mCherry-$T_{spo5}$<<$nat^r$* | Figure 4, S8-9 |
| GP9703 | *$h^{90}$ rec25-204-GFP::kanMX6 $rec8^+$-mCherry-$T_{spo5}$<<$nat^r$* | Figure 4, S8-9 |
| GP9706 | *$h^{90}$ mug20-GFP::kanMX6 $rec8^+$-mCherry-$T_{spo5}$<<$nat^r$* | Figure 4, S8-9 |
| GP9748 | *$h^-/h^-$ pat1-as1(L95G)-kanMX/ pat1-as1(L95G)-kanMX rec27-302-GFP::HygMX6/ rec27-302-GFP::HygMX6* | Figure 2, 3D-F, S6B, S7 |

| | | |
|---|---|---|
| | *rec8-mcherry-natMX/rec8-mcherry-natMX lys4-95/+ +/his4-239* | |
| GP9838 | *$h^-/h^-$ pat1-as1(L95G)-kanMX/ pat1-as1(L95G)-kanMX rec25-303-GFP::HygMX6/ rec25-303-GFP::HygMX6 rec27-238/ rec27-238 lys4-95/+ +/his4-239* | Figure S11 |
| GP9858 | *$h^-/h^-$ pat1-as1(L95G)-kanMX/ pat1-as1(L95G)-kanMX rec27-302-GFP::HygMX6/ rec27-302-GFP::HygMX6 rec12-285::natMX6/rec12-285::natMX6 lys4-95/+ +/his4-239* | Figure 3D-F |
| GP9866 | *$h^-/h^-$ pat1-as1(L95G)-kanMX/ pat1-as1(L95G)-kanMX rec27-302-GFP::HygMX6/ rec27-302-GFP::HygMX6 rec12-164(Y98F)/ rec12-164(Y98F) lys4-95/+ +/his4-239* | Figure 3D-F |
| GP9871 | *$h^-/h^-$ pat1-as1(L95G)-kanMX/ pat1-as1(L95G)-kanMX rec25-303-GFP::HygMX6/ rec25-303-GFP::HygMX6 rec27-240/ rec27-240 lys4-95/+ +/his4-239* | Figure S11 |

[a] Sources of alleles other than mating type and commonly used auxotrophies are: *ade6-3049* (Steiner and Smith, 2005); *mug20-GFP::kanMX6* (Estreicher et al.,2012); *mug20::natMX6* (Estreicher et al.,2012); *mug20-250* (Ma et al., 2017); *mug20-251* (Ma et al., 2017); *mus81::kanMX* (Smith et al., 2003); *pat1-114* (Iino and Yamamoto, 1985); *pat1-as1(L95G)* (Guerra-Moreno et al., 2012); *$rec8^+$-mCherry-$T_{spo5}$<<$nat^r$* (Ishiguro et al., 2010); *rec10-175::kanMX6* (Ellermeier and Smith, 2005); *rec10-203-GFP::kanMX6* (Fowler et al, 2013); *rec12-164(Y98F)* (Davis and Smith, 2003); *rec12-169::3HA-6His-kanMX6 [rec12Δ]* (Davis and Smith, 2003); *rec25-180::kanMX6* (Martin-Castellanos et al. 2005); *rec25-204-GFP::kanMX6* (Davis et al, 2008); *rec25-236* (Ma et al., 2017); *rec27-184::kanMX6* (Martin-Castellanos et al. 2005); *rec27-205-GFP::kanMX6* (Davis et al, 2008); *rec27-238* (Ma et al., 2017); *rec27-239* (Ma et al., 2017); *rec27-240* (Ma et al., 2017); *red50S* (Farah et al., 2002).

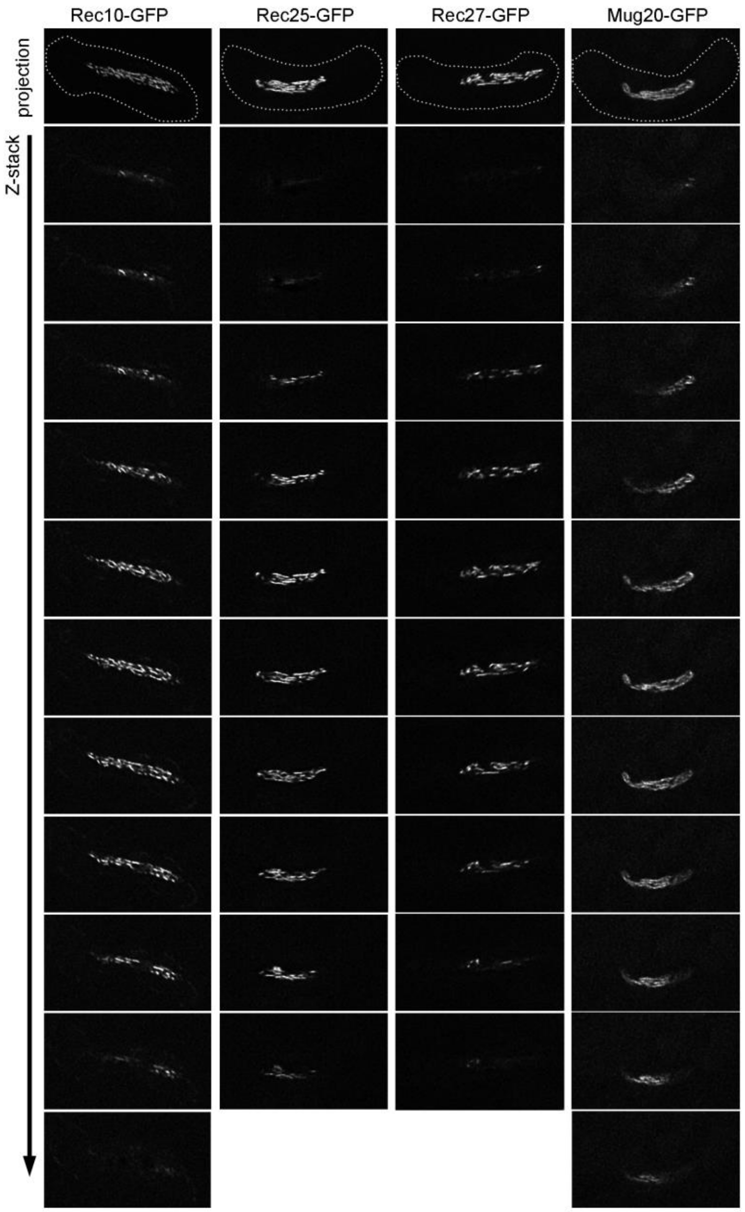


**Figure S1. LinE subunits form linear structures in zygotic meiosis at 25°C**

Rec10-GFP, Rec25-GFP, Rec27-GFP and Mug20-GFP were observed in $h^{90}$ strains using Structured Illumination Microscopy (SIM). Each strain was incubated on sporulation medium (MEA) at 25°C for 13 – 15 hr. Each image is representative of at least 20 cells in the horsetail stage (*i.e.*, meiotic prophase; Ding et al., 2004). All four LinE subunits formed continuous, linear structures. The first image of each strain shows the maximal projection of the entire Z-stack of image sections, shown consecutively below. The dotted line represents the outline of the cell. See also Figure 1A.

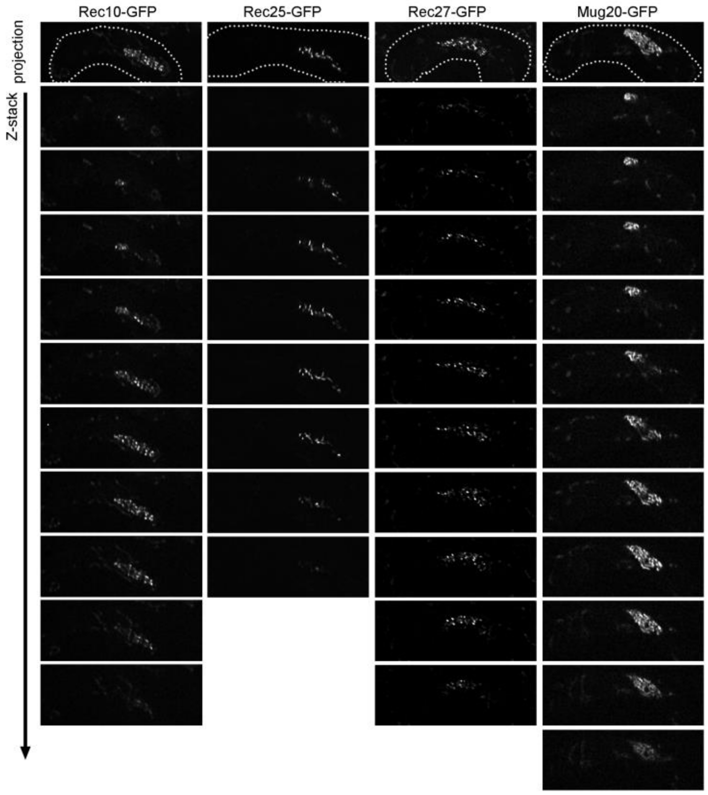


**Figure S2. Rec10-, Rec27- and Rec25-GFP fail to form linear structures at high temperature (34°C) in zygotic meiosis**

Rec10-GFP, Rec25-GFP, Rec27-GFP and Mug20-GFP were observed in $h^{90}$ strains using Structured Illumination Microscopy (SIM). Each strain was incubated on sporulation medium (MEA) at 25°C for 13 – 15 hr. Each image is representative of at least 20 cells in the horsetail stage (*i.e.*, meiotic prophase; Ding et al., 2004). The morphologies of Rec10-, Rec-25- and Rec27-GFP were non-continuous and often showed bright nuclear dots. These "dotty foci" were different from the "linear structures" observed at 25°C (Figures 1 and S1) but were similar to the structures in previous studies of azygotic meiosis at 34°C (Lorenz et al., 2004;Davis et al., 2008; Estreicher et al., 2012; Fowler et al., 2013). Mug20-GFP showed long linear structures at 34°C, similar to those at 25°C (Figure 1). The first image of each strain shows the maximal projection of the entire Z-stack of image sections, shown consecutively below. The dotted line represents the outline of the cell. See also Figure 1B.

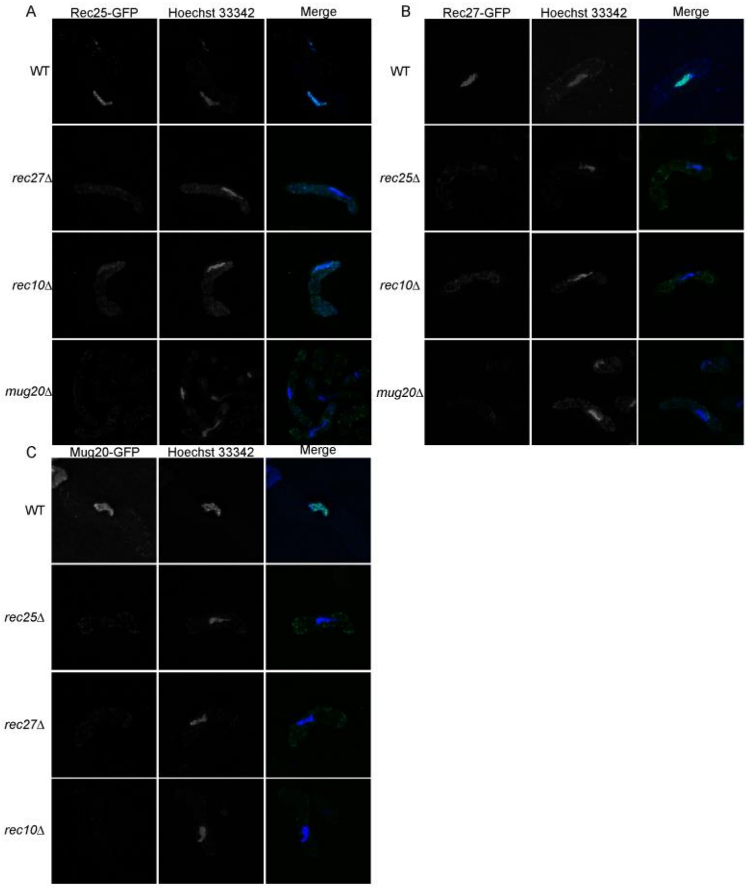


**Figure S3. Rec25-, Rec27-, and Mug20-GFP nuclear structures are interdependent with other LinE subunits**

**(A)** Rec25-GFP, **(B)** Rec27-GFP and **(C)** Mug20-GFP were observed in the indicated LinE subunit mutants in $h^{90}$ cells using a high-resolution microscope (DeltaVision). Cells were incubated on MEA at 25°C for 14 – 16 hr. Each strain was incubated on sporulation medium (MEA) at 25°C for 13 – 15 hr. Each image is representative of at least 20 cells in the horsetail stage (*i.e.*, meiotic prophase; Ding et al., 2004). Nuclear localization is shown by Hoechst 33342 staining.

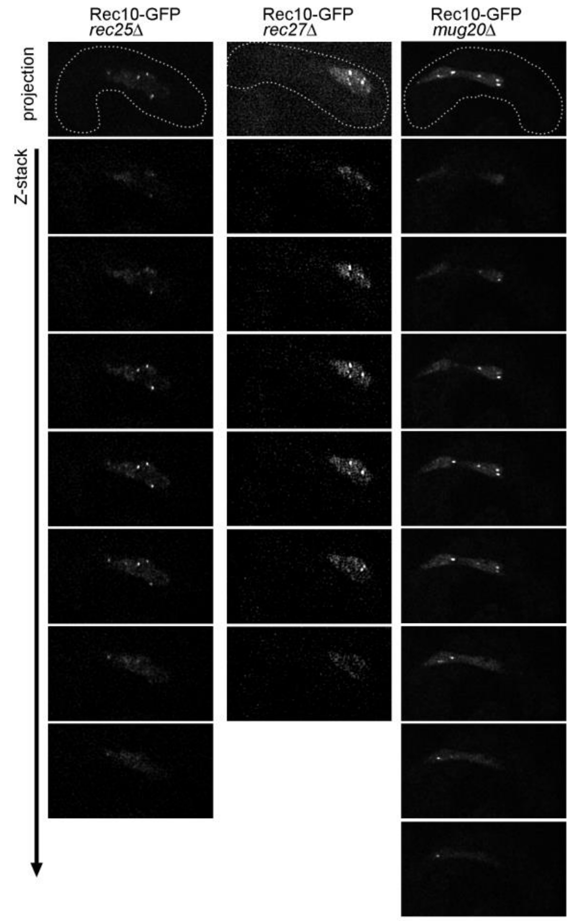


**Figure S4. Rec10-GFP enters the nucleus and forms nuclear foci independent of other LinE subunits at 25°C in zygotic meiosis**

Morphology of Rec10-GFP was observed in $h^{90}$ strains with *rec25Δ, rec27Δ* or *mug20Δ* genetic background using Structured Illumination Microscopy (SIM). Cells were incubated on MEA at 25°C for 13 – 15 hr. Each strain was incubated on sporulation medium (MEA) at 25°C for 13 – 15 hr. Each image is representative of at least 20 cells in the horsetail stage (*i.e.*, meiotic prophase; Ding et al., 2004). Rec10-GFP showed uniform nuclear distribution in all three LinE subunit mutants. A few bright Rec10-GFP foci were often found in the nucleus. These results indicate that Rec10-GFP still enters the nucleus and forms nuclear foci, but not linear structures, in the absence of any other LinE subunit. The first image of each strain shows the maximal projection of the entire Z-stack of image sections, shown consecutively below. The dotted line represents the outline of the cell. See also Figure 1C.

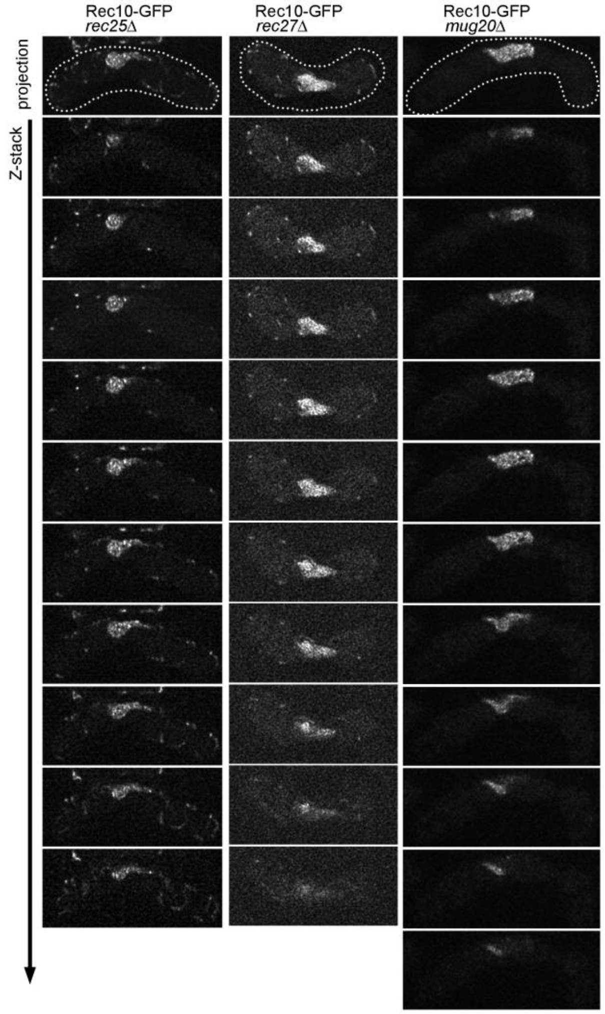


**Figure S5. Rec10-GFP shows similar morphology at 34°C and 25°C in zygotic meiosis when other LinE subunits are missing**

Rec10-GFP was observed in *h*$^{90}$ strains with *rec25Δ, rec27Δ* or *mug20Δ* genetic background using Structured Illumination Microscopy (SIM). Cells were incubated on MEA at 34°C for 10 – 17 hr. Each image is representative of at least 20 cells in the horsetail stage (*i.e.*, meiotic prophase; (Ding et al., 2004). At both 25°C and 34 °C (see also Figures 1A and 1B), Rec10-GFP showed uniform nuclear distribution with a few bright nuclear foci in the absence of any other LinE subunit. Thus, other LinE subunits are more critical than temperature for Rec10-GFP structure formation. The first image of each strain shows the maximal projection of the entire Z-stack of image sections, shown consecutively below. The dotted line represents the outline of the cell. See also Figure 1D.

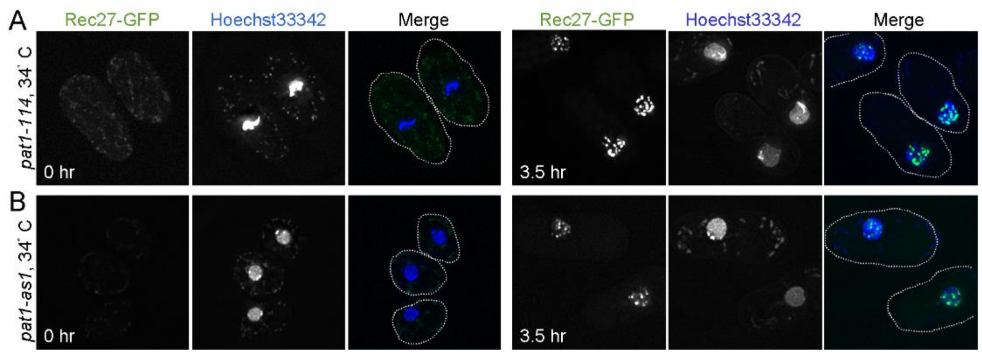


**Figure S6. Rec27-GFP forms "dotty" foci in synchronized azygotic meiosis at 34°C**

Rec27-GFP was observed in live cells of synchronized, azygotic meiosis cultures (*pat1-114* or *pat1-as1*) at 34°C. Cells were collected at the indicated times after meiotic induction and were observed with high-resolution fluorescence microscopy (DeltaVision). Each image is representative of at least 20 cells. Rec27-GFP showed "dotty" nuclear foci at 3.5 hr after meiotic induction in **(A)** *pat1-114* and **(B)** *pat1-as1* strains at 34°C. As in Figure 2, Rec27-GFP showed bright, non-continuous nuclear "dots," consistent with previous observations under similar conditions (Lorenz et al., 2004; Davis et al., 2008; Estreicher et al., 2012; Fowler et al., 2013). Nuclear localization is shown by Hoechst 33342 staining. The dotted line represents the outline of the cell.

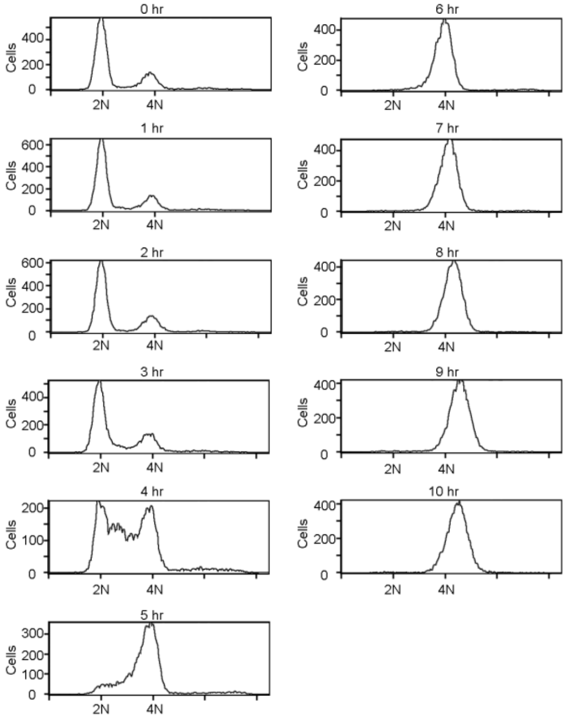


**Figure S7. Meiotic DNA replication in azygotic meiosis (*pat1-as1*) at 25°C**

DNA content of azygotic meiosis time-course samples were analyzed by flow cytometry. The 0 hr sample was used for identifying DNA content (2N or 4N). Y axis indicates cell counts.

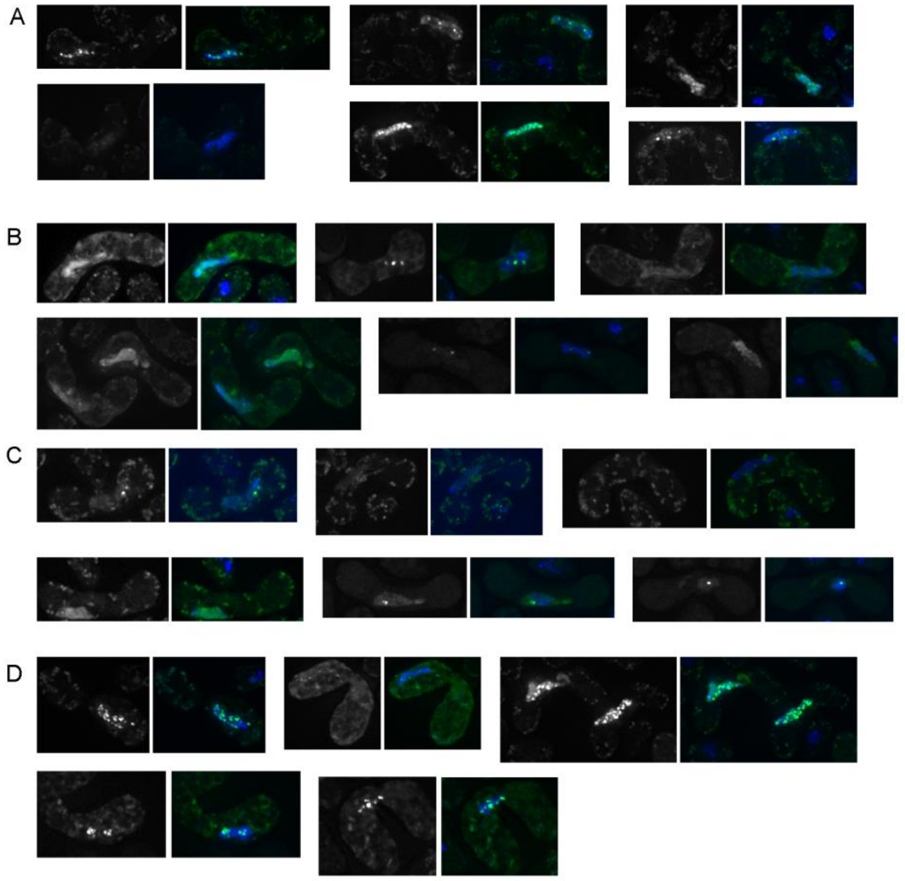


**Figure S8. LinE subunits are sensitive to 1,6-hexanediol treatment**

More examples of **(A)** Rec10-GFP, **(B)** Rec25-GFP, **(C)** Rec27-GFP and **(D)** Mug20-GFP in *h*$^{90}$ strains after 1,6-hexanediol treatment. Cells were incubated at 25°C for 14 – 16 hr and observed in high-resolution fluorescence microscopy (DeltaVision). 1,6-Hexanediol (10% in EMM2-N) was added for five min at room temperature before observation. Nuclear localization is shown by Hoechst 33342 staining. In each pair of panels, the left panel shows LinE-GFP, and the right panel shows merged images of LinE-GFP and Hoechst 33342 staining.

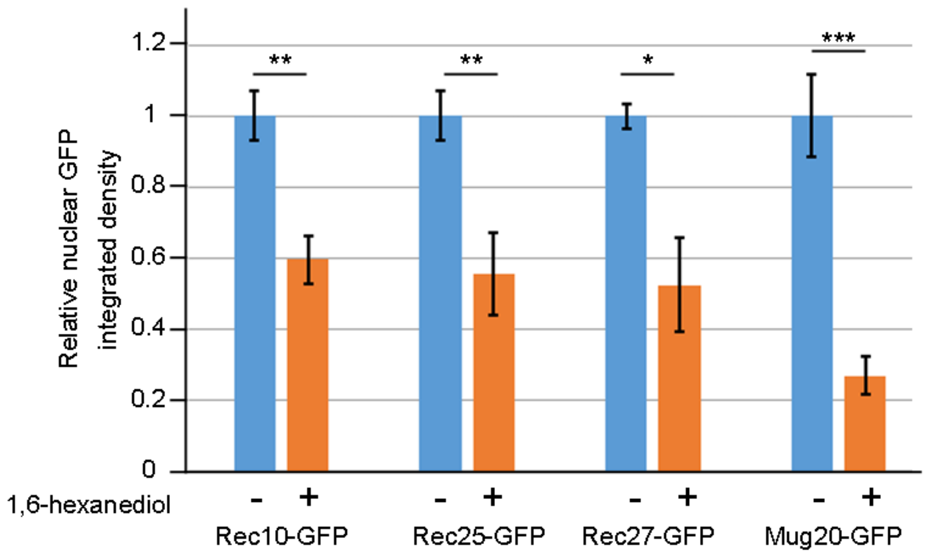


**Figure S9. Nuclear LinE-GFPs are reduced by 1,6-hexanediol**

Nuclear Rec10-GFP, Rec25-GFP, Rec27-GFP and Mug20-GFP were quantified in $h^{90}$ strains with or without 1,6-hexanediol treatment. Y axis indicates the mean value of the nuclear GFP integrated density relative to that of the untreated samples. At least six cells were quantified in each treatment. The error bar indicates standard error of the mean (SEM); *(P < 0.05), **(P < 0.01), and ***(P < 0.001) indicate significance of the difference from wild type by unpaired t-test.

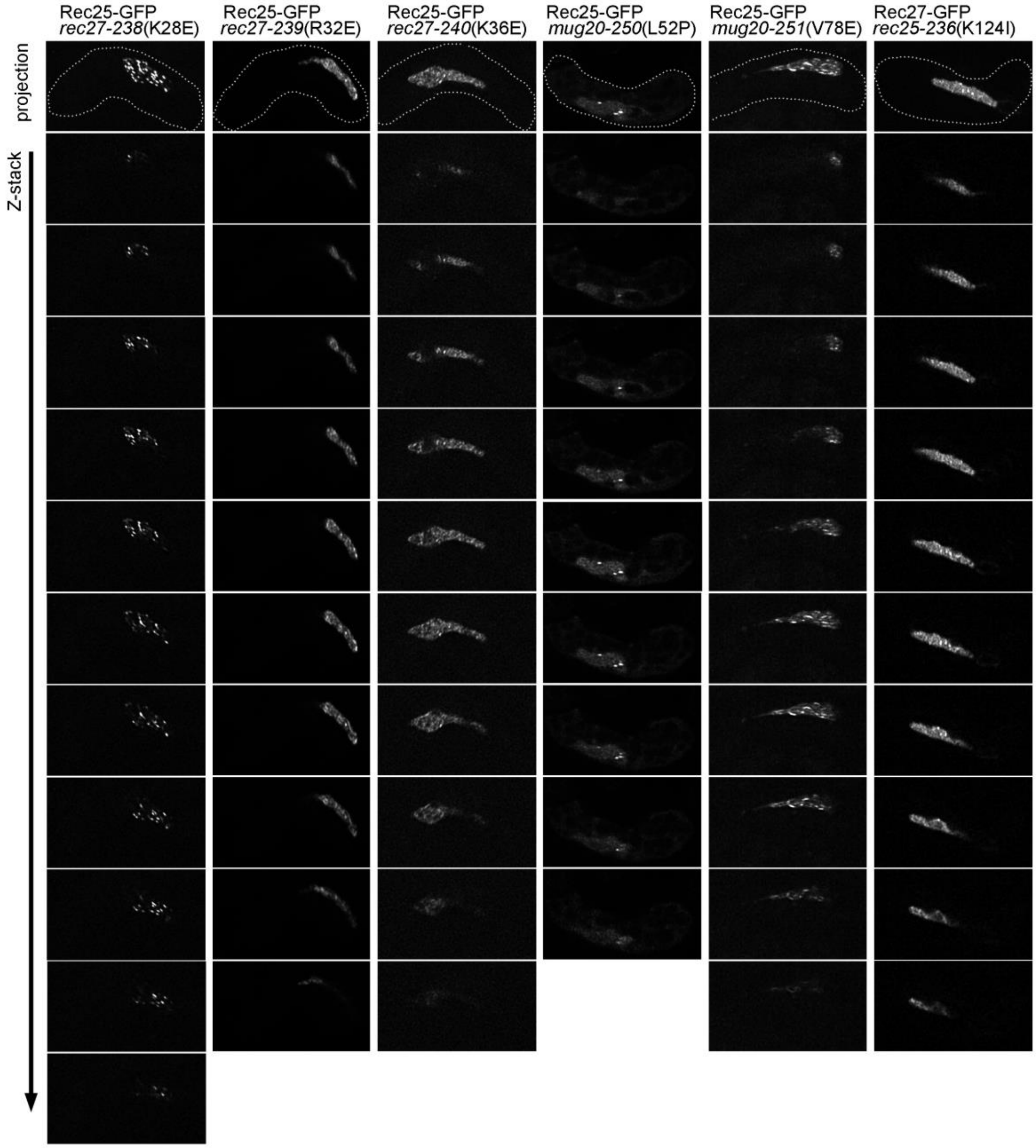


**Figure S10. LinE missense mutations alter LinE linear structures**

Rec25-GFP (in *rec27-238*, *rec27-239*, *rec27-240*, *mug20-250* and *mug20-251*) and Rec27-GFP (in *rec25-236*) were observed in the indicated LinE subunit mutants in $h^{90}$ strains using Structured Illumination Microscopy (SIM). Cells were incubated on MEA at 25°C for 14 – 16 hr. Each image is representative of at least 20 cells in the horsetail stage (*i.e.*, meiotic prophase; Ding et al., 2004). The first image of each strain shows the maximal projection of the entire Z-stack of image sections, shown consecutively below. The dotted line represents the outline of the cell. See also Figure 5.

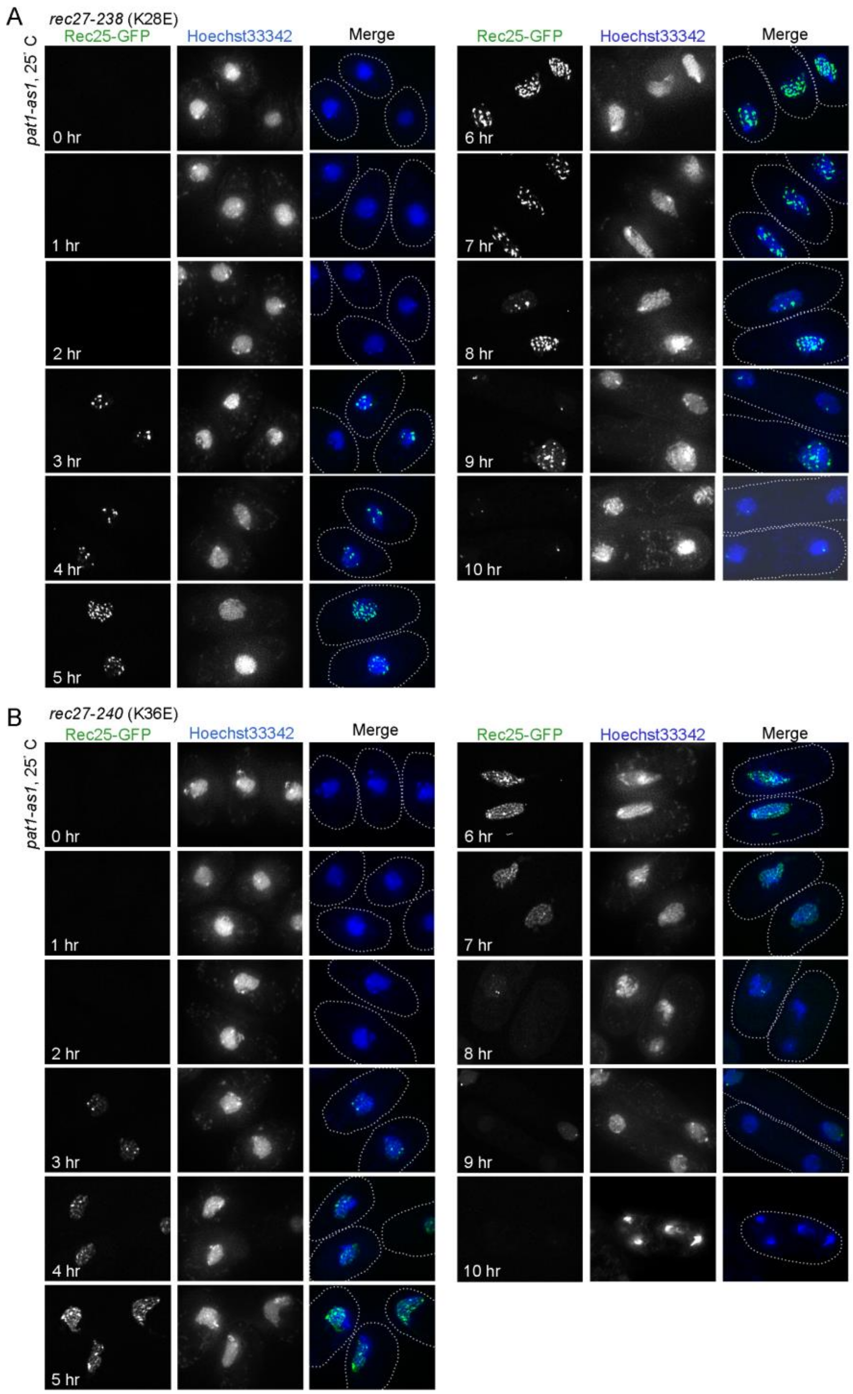


**Figure S11. LinE missense mutations alter LinE linear structure**

Rec25-GFP in **(A)** *rec27-238*(K28E) or **(B)** *rec27-240*(K38E) was observed in live cells of synchronized, azygotic meiotic cultures (*pat1-as1*) at 25°C. Cells were collected at the indicated times after meiotic induction and were observed with Structured Illumination Microscopy (SIM). Each image is representative of at least 20 cells analyzed in each condition. Rec25-GFP never formed linear configuration in either missense mutant during meiotic induction. Nuclear location is shown by Hoechst 33342 staining. The dotted line represents the outline of the cell.
